# A Genetically Engineered Primary Human Natural Killer Cell Platform for Cancer Immunotherapy

**DOI:** 10.1101/430553

**Authors:** Emily J. Pomeroy, John T. Hunzeker, Mitchell T. Kluesner, Margaret R. Crosby, Walker S. Lahr, Laura Bendzick, Jeffrey S. Miller, Beau R. Webber, Melissa A. Geller, Bruce Walcheck, Martin Felices, Timothy K. Starr, Branden S. Moriarity

## Abstract

Tumors can evade natural killer (NK) cells by activating inhibitory pathways. We therefore have developed a highly efficient CRISPR/Cas9-based method for editing the genome of peripheral blood human NK cells (PB-NKs) to knock out ADAM17 and PD1 or knock-in genes using recombinant AAV6. Our method allows editing of PB-NKs at efficiencies reaching 90%, equivalent to methods reported for primary human T cells. Moreover, we demonstrate that ADAM17 and PD1 KO PB-NKs have significantly improved activity, cytokine production, and cancer cell cytotoxicity. Our platform represents a feasible method for generating engineered NK cells as a universal therapeutic for cancer immunotherapy.

## Introduction

Natural killer (NK) cells are critical components of the innate immune system due to their ability to kill a variety of target cells, including cancer cells. Killing of targets by NK cells is mediated by the integration of signals from activating and inhibitory receptors and cytokines^1^. The ability of NK cells to kill tumor cells, along with the ease with which they are isolated and expanded from peripheral blood, has made them an attractive source for immunotherapy^2,3^. Moreover, as they do not induce graft versus host disease^4^, they can be generated from unrelated donors and, therefore, represent an ‘off the shelf’ therapeutic cell type. However, NK cell immunotherapy has seen limited success in the clinic, due in part to their lack of persistence and expansion after transplant^5^. Changes in NK cell receptor repertoire and ligand expression in the tumor microenvironment can also lead to decreased NK cell activity^6,7^.

One potent activator of NK cells is CD16a (FCγRIIIa), which binds the Fc portion of IgG-coated target cells. This antibody-dependent cell-mediated cytotoxicity (ADCC) by NK cells underlies the effectiveness of anti-tumor therapeutic antibody treatment of malignancies^8,9^. After activation, CD16a is rapidly cleaved from the cell surface by A Disintegrin And Metalloproteinase-17 (ADAM17), resulting in attenuated activity. Chemical inhibition of ADAM17 leads to increased cytokine production by human NK cells^10^, suggesting that the use of monoclonal antibodies combined with inhibition of ADAM17 could enhance the anti-tumor response.

Various cells within the tumor micro-environment also secrete immunosuppressive molecules that interfere with the complex array of receptors that regulate NK cells and their ability to expand^12^. For example, some tumor cells upregulate PD-L1, which engages PD1 on NK cells reducing cytotoxic activity^13,14^. This inhibition imposed on NK cells is a limiting factor in effective NK immunotherapy and overcoming this immunosuppression is necessary for viable therapeutic development. Strategies for overcoming these challenges have focused on releasing NK cell inhibition through the use of pharmaceutical inhibitors and monoclonal antibodies^15–17^. However, there are limitations to these strategies: off-target toxicity is common with these types of therapies and, importantly, no humanized antibodies or chemical inhibitors exist for some high interest targets.

An alternative approach is to use CRISPR/Cas9 to edit genes relevant to NK function in order to improve their utility as an immunotherapeutic agent. Changes made with this approach are confined to NK cells, reducing undesirable systemic effects. Additionally, design and optimization of CRISPR guide RNAs (gRNAs) is straightforward and cost-effective, making it easy to target any gene of interest. Here, we present the development of an approach for achieving high rates of gene editing in activated primary human NK cells combined with expansion to clinically relevant numbers and show that gene edited NK cells have enhanced function *in vitro* and *in vivo*. Our method is capable of knocking-out genes at rates up to 90% and knocking in genes at rates up 79%. In summary, we present a rapid platform for generating high functioning genetically modified NK cells for use in cancer immunotherapy.

## Materials and Methods

### Donor NK Cell Isolation and Expansion

Peripheral blood mononuclear cells (PBMCs) from de-identified healthy human donors were obtained by automated leukapheresis (Memorial Blood Centers, Minneapolis, MN) and further isolated on a ficoll-hypaque (Lonza) gradient. CD56+CD3− NK cells were isolated by negative selection using the EasySep Human NK Cell Enrichment Kit (Stemcell Technologies). After isolation, NK cells were either rested by culture with 1 ng/mL IL15, or activated by co-culture with Clone9.mbIL21 at a 2:1 or 1:1 (feeder:NK) ratio in B0 supplemented with 50 IU/mL IL2 (Peprotech), as described^18^. Samples were obtained after informed consent with approval from the University of Minnesota Institutional Review Board (IRB 1602E84302).

### Cell Culture

NK cells were maintained in B0 medium^19^ supplemented with 1 ng/mL IL15 unless otherwise noted. NK cells were cryopreserved at 1×10^7^ cells/mL in CryoStor CS10 (Sigma Aldrich). The transgenic Clone9.mbIL21 cell line, the human Burkitt’s lymphoma cell line Raji, the human erythroleukemia cell line K562, the human prostate cancer cell line DU145, and the human monocytic leukemia cell line THP1 were maintained in RPMI 1640 (GE Healthcare Life Sciences) supplemented with 10% FBS (GE Healthcare Life Sciences) and 100 U/mL penicillin (Corning) and 100 U/mL streptomycin (Corning). The human ovarian cancer cell line MA148 was maintained in DMEM (GE Healthcare Life Sciences) supplemented with 10% FBS and 100 U/mL penicillin and 100 U/mL streptomycin.

### Guide RNA design

gRNAs targeting ADAM17, PD1, CISH, and AAVS1 were designed using the CRISPR MIT webtool (http://crispr.mit.edu.ezp1.lib.umn.edu/) along with Cas-OFFinder. Six gRNAs per target gene were tested in HEK293T cells and the most efficient gRNA (see Table 1) was then ordered from TriLink Biotechnologies or Synthego with 2’-O-methyl and 3’ phosphorothioate modifications to the first three 5’ and the last three 3’ nucleotides.

**Table 1.**
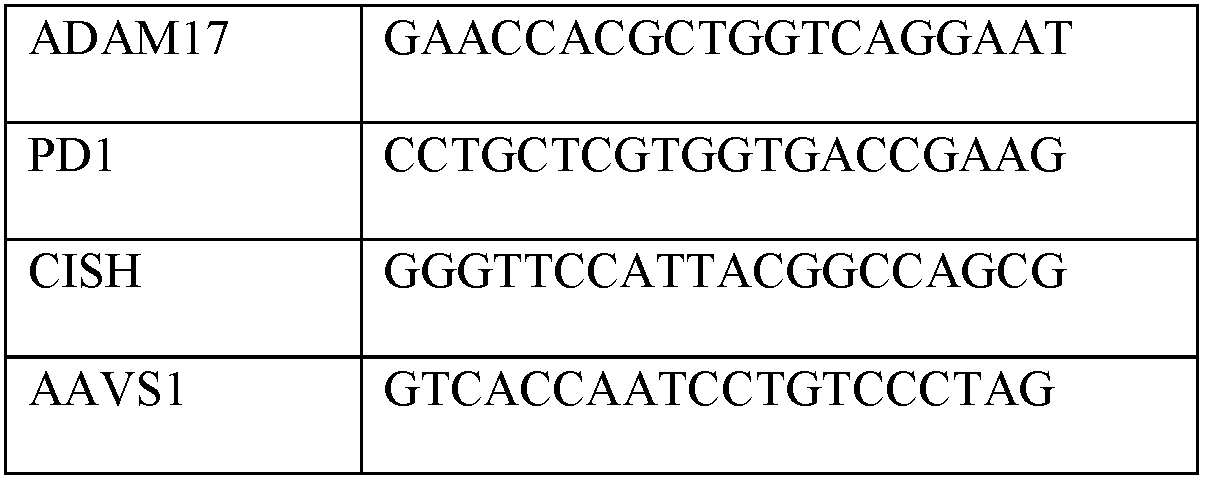
Guide RNA target sequences.

### Electroporation of activated NK cells

Activated NK cells were pelleted and resuspended at 3×10^7^ cells/mL in T buffer (Neon Transfection System Kit, ThermoFisher Scientific). 1.5 ug Cas9 mRNA (Trilink) and 1 ug chemically-modified guide RNA (Trilink or Synthego) were added to 10 uL (3×10^5^ cells) on ice. Cas9 mRNA alone, without guide RNA, was used as a control for all experiments. This mixture was electroporated with the Neon Transfection System (ThermoFisher Scientific) using 2 pulses of 1850 volts and 10 ms pulse width. NK cells were recovered in warm B0 medium containing 1 ng/mL IL15. For rAAV6 infection, rAAV6 (MOI=500,000) was added to NK cells 30 minutes after electroporation. rAAV6 particles were produced by the University of Minnesota Vector Core.

### Analysis of gene editing efficiency

PCR primers were designed to amplify a 400–500 bp region surrounding the gRNA target site (Table 2). Five days after electroporation, genomic DNA was PCR amplified in one step using AccuPrime Taq DNA Polymerase (Invitrogen). For analysis by TIDE, PCR amplicons were Sanger sequenced (ACGTinc or University of Minnesota Genomics Center) and Sanger chromatograms were uploaded to the TIDE webtool (https://tide-calculator.nki.nl). For Next Generation Sequencing (NGS), primers with Nextera universal primer adaptors (Illumina) were designed to amplify a 350–475 bp site surrounding the region of interest (Table 2). Samples were submitted to the University of Minnesota Genomics Center for subsequent amplification with indexing primers and sequencing on a MiSeq 2×200 bp run (Illumina). A minimum of 1000 read-pairs were generated per sample. Raw .fastq files were analyzed against a reference sequence and gRNA protospacer sequence using the CRISPR/Cas9 editing analysis pipeline CRISPR-DAV, as previously described^20^.

**Table 2.**
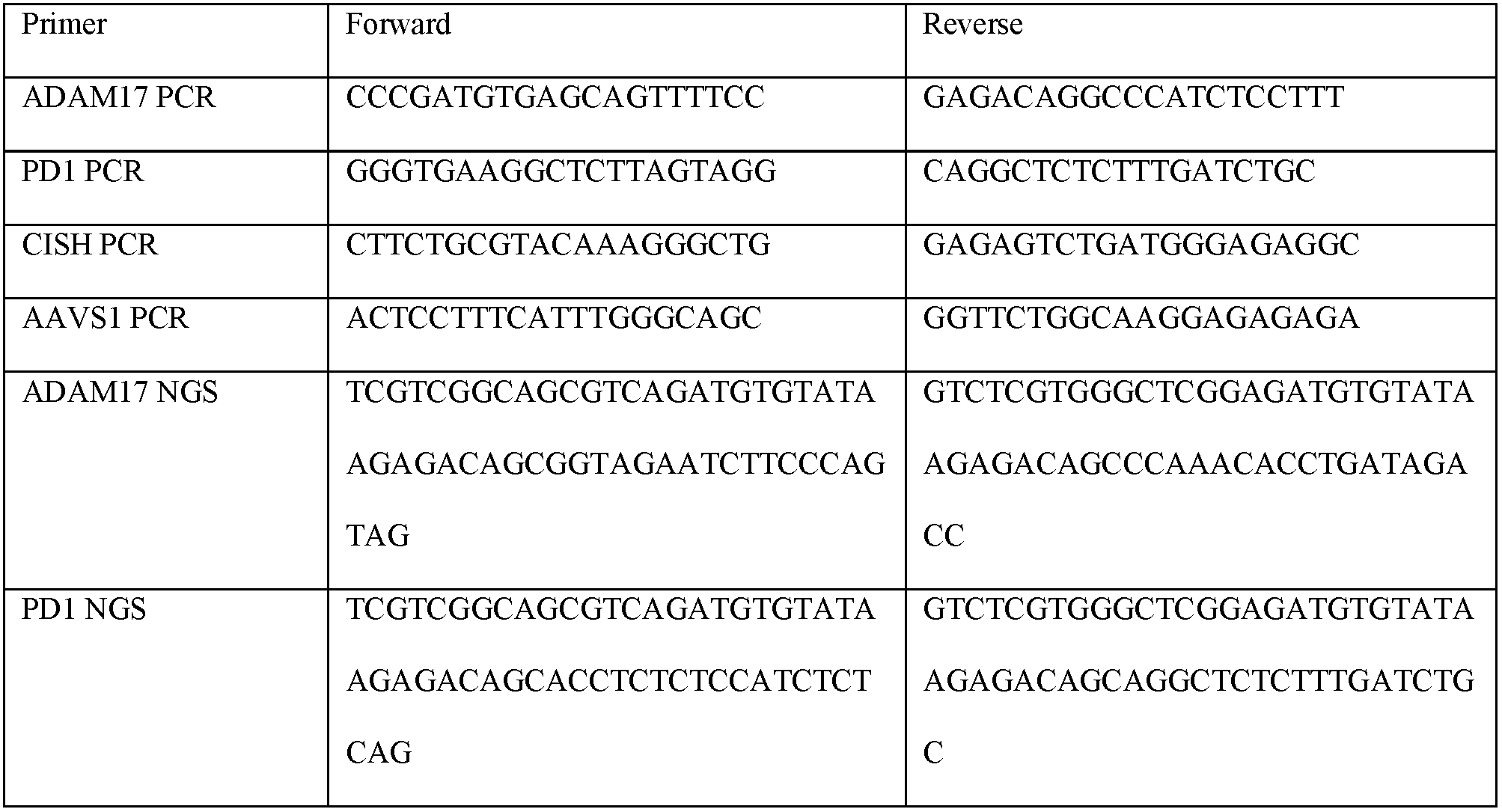
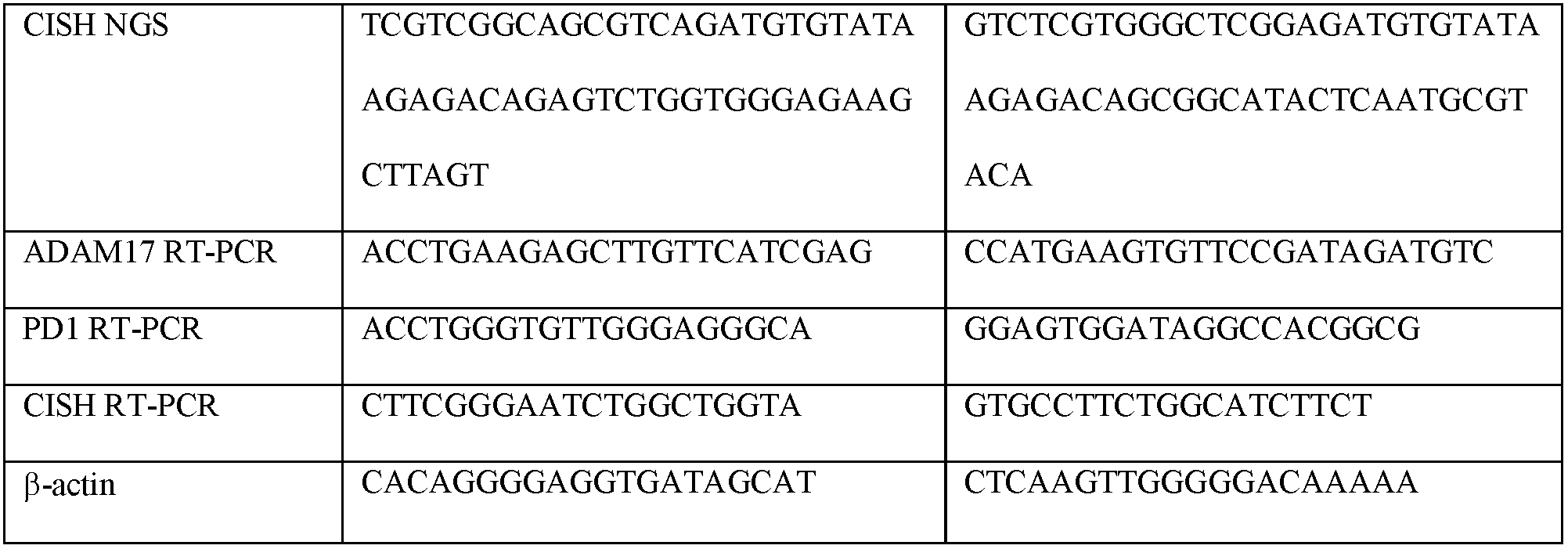
Primer sequences.

### RT-PCR

RNA was extracted using the PureLink RNA Mini Kit (Ambion). cDNA was generated using the Transcriptor First Strand cDNA Synthesis Kit (Roche). Real-time PCR was conducted using intron-spanning primer sets (Table 2) and SYBR Green Master Mix (Applied Biosystems) and analyzed using a CFX-96 (Bio-Rad). Gene expression levels were calculated relative to β-actin.

### Western blot

Cells were lysed in RIPA buffer (Sigma) supplemented with phosphatase inhibitors (Sigma) and protease inhibitors (Roche) for 30 min and purified by centrifugation (14,000 × *g* for 15 min). Concentration of the protein lysate was determined by BCA (Pierce). All samples and reagents were prepared according to the SimpleWestern manual for Wes (ProteinSimple). Samples were diluted to equal concentration (750 ng) with 0.1X Sample Buffer (ProteinSimple) and combined with 0.2 mg/mL 5X Fluorescent Master Mix (ProteinSimple). Final samples were denatured at 95ºC for 5 minutes. The provided microplate containing separation and stacking matrices was loaded with biotinylated ladder (ProteinSimple), blocking reagent, primary antibody (B-actin (Cell Signaling Technologies) 1:100; CISH (Cell Signaling Technologies) 1:50; antibodies were diluted using provided diluent), Secondary HRP conjugate, chemiluminescent luminol-peroxide mix, and wash buffer. The self-regulating protocol of the SimpleWestern system was followed and data was analyzed using the Compass Software (ProteinSimple).

### Antibodies and Flow Cytometry

The following antibodies were used: APC-, FITC-, PE-, or Biotin-conjugated anti-CD56 (clone REA196; Miltenyi Biotec), PE-conjugated anti-CD3 (clone SK7; BD Biosciences), PE/Cy7-conjugated anti-CD16A (clone 3G8; BioLegend), Brilliant violet 421-conjugated anti-CD16A (clone 3G8; BioLegend), Brilliant violet 421-conjugated anti-g (clone 4S.B3; BioLegend), FITC-conjugated anti-CD107a (clone H4A3; BD Biosciences), Brilliant violet 605-conjugated anti-CD62L (clone Dreg56; BD Biosciences), PE-conjugated anti-active caspase-3 (clone C92–605; BD Biosciences), SYTOX Blue dead cell stain (ThermoFisher), Fixable viability dye eFluor 780 (eBioscience); anti-CISH (clone D4D9; Cell Signaling Technology). CellTrace CFSE (ThermoFisher Scientific) was used to label target cells and measure proliferation. Flow cytometry assays were performed on LSRII or LSR Fortessa flow cytometers (BD Biosciences) and all data were analyzed with FlowJo version 10.4 software (FlowJo LLC).

### NK Cell Functional Assays

For PMA stimulation, NK cells were pre-treated for 1 hour with 10 uM ADAM17 inhibitor (INCB007839) or DMSO control. NK cells were then stimulated with 1 ug/mL PMA for 1 hour and CD16a expression was measured by flow cytometry. For intracellular cytokine staining, NK cells were plated at 2.5×10^5^ cells per 100 uL B0 with no cytokines added. After incubation overnight, target cells were added at the indicated E:T ratios (2:1 for assays with ADAM17 KO NK cells, 1:1 for assays with PD1 or CISH KO NK cells). For ADCC assays, Raji cells were pre-coated with Rituximab (Genentech) at 10 ug/mL for 30 minutes. FITC-conjugated anti-CD107a was added to the culture and cells were incubated for 1 hour at 37 C. Brefeldin A and monensin (BD Biosciences) were added after 1 hour and cells were incubated for an additional 5 hours. Cells were stained with fixable viability dye and for extracellular antigens and then were fixed and permeabilized using BD Cytofix/Cytoperm (BD Biosciences). Cells were then stained for intracellular IFNγ.

### Target Cell Killing Assays

NK cells were plated at 2.5×10^5^ cells per 100 uL B0 medium with no cytokines added and incubated overnight. Target cells were pelleted and labeled with CellTrace CFSE (ThermoFisher Scientific) for 10 minutes at room temperature, then washed in 10 mL FBS. For ADCC assays, CFSE-labeled Raji cells were pre-coated with Rituximab (Genentech) at 10 ug/mL for 30 minutes. CFSE-labeled target cells were added to NK cells (E:T=2:1 for ADAM17 assays, E:T=1:1 for PD1 and CISH assays). Co-cultures were incubated at 37C for 6 hours. Cells were stained with fixable viability dye and then were fixed and permeabilized with ice cold 70% ethanol for 30 minutes. Cells were then stained for active caspase-3.

### In vivo tumor model

All animal studies were approved by the University of Minnesota Institutional Animal Care and Use Committee (IACUC 1610–34201A). NOD/SCID/γc^−/−^(NSG) mice were purchased from Jackson Laboratories and used for all *in vivo* experiments. Mice were given 2×10^5^ luciferase-expressing MA148 cells via IP injection four days prior to NK cell delivery (day −4). On day −1, mice were sub-lethally irradiated (225 cGy), and tumor burden was measured using bioluminescent imaging (BLI). Mice were then grouped based on BLI to ensure each treatment group started with the same average tumor burden. On day 0, NK cells (1×10^6^ cells per mouse) were delivered IP. BLI was measured weekly using the Xenogen IVIS 50 Imaging System (Caliper Life Science). Mice received IP injections of IL-15 (5 ug/mouse) every Monday, Wednesday, and Friday for 3 weeks. Animal health was monitored daily and mice were euthanized when moribund.

### Statistical Analysis

The Student’s t test was used to test for significant differences between two groups. Differences between more than 2 groups were tested by one-way ANOVA analysis with Tukey post-hoc test for multiple comparisons. All in vitro assays were repeated in 3–5 independent donors. Mean values +/− SEM are shown. In vivo data was generated using n=10 mice per group. Overall survival was calculated using Kaplan-Meier methods and treatment groups were compared using log rank tests. P values < 0.05 were considered statistically significant. Statistical analyses were performed using GraphPad Prism 6.0.

## Results and Discussion

To improve NK cell cytotoxicity, we developed an optimized CRISPR/Cas9 system capable of disrupting important regulatory genes in activated primary human NK Cells. Based on our previous work with primary human T cells and B cells, we hypothesized that activating NK cells prior to electroporation would make them more amenable to nucleic acid delivery^21,22^. To test this, we delivered nucleic acid in the form of eGFP mRNA to NK cells isolated from peripheral blood mononuclear cells (PBMCs) using protocols with resting or activated NK cells. We found that NK cells activated with irradiated feeder cells expressing membrane-bound IL21 (Clone9.mbIL21, C9s)^18^ and cultured with IL-2 for 7 days prior to electroporation resulted in the most efficient nucleic acid delivery and NK cell viability (**Figure 1A**). Using this approach, we routinely achieve >98% transfection efficiency with >90% cell viability (**Supplemental Figure 1A**). Importantly, C9-expanded NK cells have been used clinically and have been shown to be safe and effective for multiple myeloma patients^23^.

**Figure 1.**
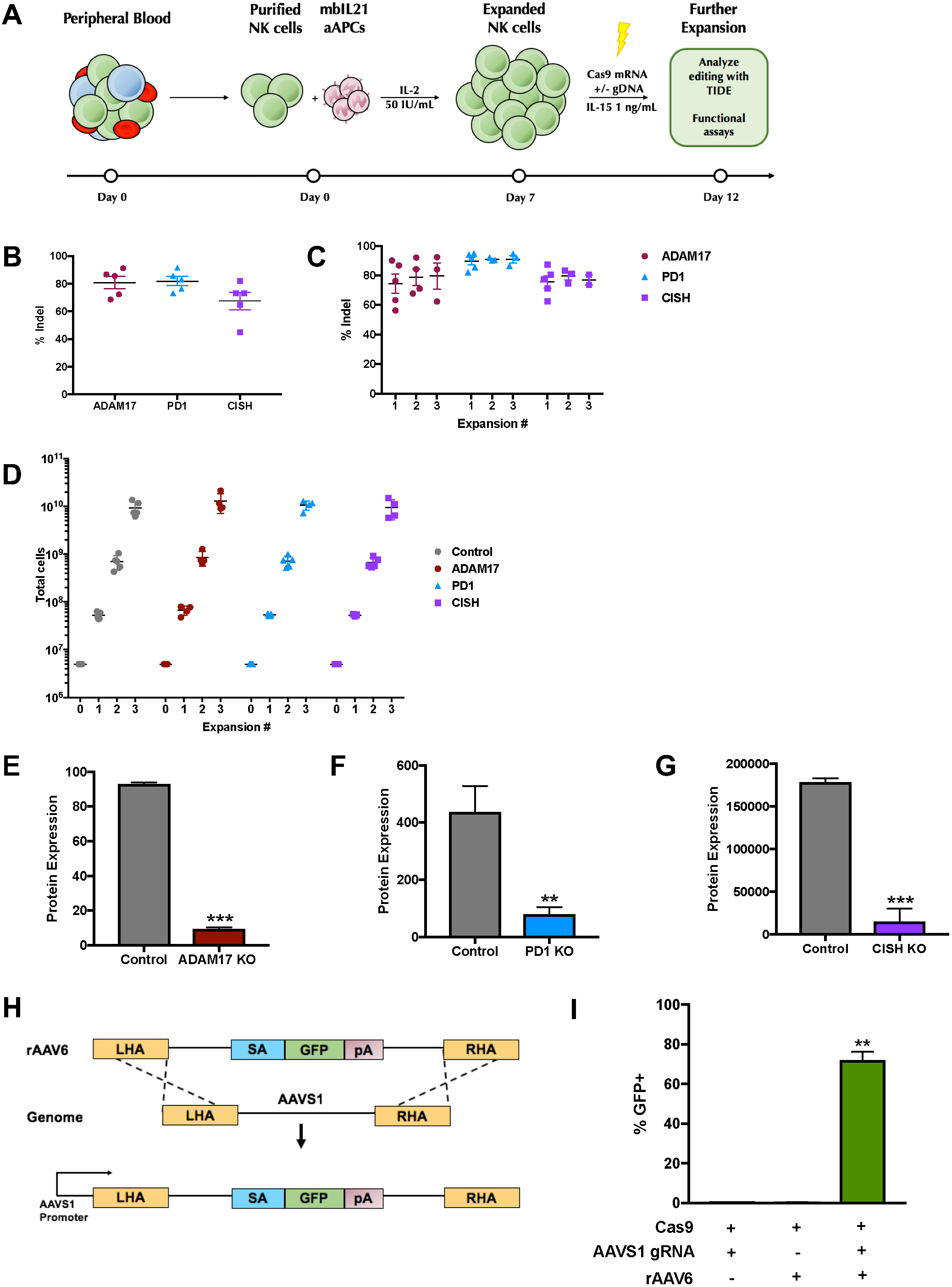
A highly efficient method of expansion and CRISPR/Cas9-based engineering of primary human NK cells. **(A)** Protocol for expansion and gene editing of primary human NK cells. **(B)** Next generation sequencing (NGS) analysis of indel formation after delivery of Cas9 mRNA + gRNA targeting ADAM17 (maroon), PD1 (blue), and CISH (purple) (n=5 independent donors). **(C)** Analysis of indel formation by TIDE over serial expansions with Clone9.mbIL21 feeder cells (n=5 independent donors). **(D)** Expansion of engineered cells to clinically relevant numbers using multiple rounds of Clone9.mbIL21 expansion (n=5 independent donors). **(E-G)** Indel formation in target genes leads to protein knockout. Flow cytometry analysis of % ADAM17+ NK cells (**E**, n=2 independent donors, ***P=0.0002, Student’s t test), PD1 mean fluorescence intensity in NK cells (**F**, n=4 independent donors, **P=0.0089, Student’s t test) expression, and ProteinSimple Wes analysis of CISH expression relative to β-actin (**G**, n=3 independent donors, ***P=0.0005, Student’s t test) after gene knockout. **(H)** Gene knock-in strategy to integrate a splice-acceptor-EGFP cassette to the AAVS1 locus using rAAV6 and CRISPR/Cas9. **(I)** EGFP expression 14 days after gene knock-in with rAAV6 (n=2 independent donors, **P=0.0016, Student’s t test)

Using our optimized protocol we delivered Cas9 mRNA alone as a control, or in combination with synthetic chemically modified gRNAs^24^ targeting *ADAM17*^11^, *PD1*^14^*, and CISH*^25^ to C9 activated NK cells. We were able to achieve gene editing efficiency of 80.9 ± 9.9% for *ADAM17*, 81.9 ± 7.4% for *PD1*, and 67.6 ± 14.1% for *CISH* without the use of any enrichment methods. (**Figure 1B**). We compared next generation sequencing (NGS) and Tracking of Indels by Decomposition (TIDE) analysis to quantify gene editing efficiency across five independent donors in which we knocked out *ADAM17*, *PD1*, or *CISH*. The TIDE web tool quantifies indel formation using Sanger sequencing reactions^26^. We observed no significant difference between editing efficiency calculated using TIDE versus NGS (**Supplemental Figure 1G**). Critically, these editing events were maintained at similar frequencies after multiple rounds of expansion using C9 feeders (**Figure 1C**) and did not affect our ability to expand cells to clinically relevant numbers (**Figure 1D**). Protein expression of targeted genes was also significantly decreased by 89.8 ± 1.2% for ADAM17, 82.7 ± 8.6% for PD1, and 91.9 ± 14.0% for CISH, and mRNA expression followed a similar pattern (**Figure 1E-G; Supplemental Figure 1B-F**). Furthermore, with the goal of developing “off-the-shelf” clinical products we sought to optimize cryopreservation of activated and gene-edited NK cells. Previous groups have shown low NK cell recovery and cytotoxicity after cryopreservation^27^. We found that freezing 1×10^7^ NK cells per mL using CryoStor CS10 preservation media yielded ~80% recovery after thaw, and that gene editing did not affect this process (**Supplemental Figure 1H**).

In addition to successful gene knock-out (KO), we adapted our method for gene knock-in (KI) by co-delivering a DNA template for homologous recombination (HR) using rAAV6 along with Cas9 and gRNA. As proof-of-principle, we delivered Cas9 and gRNA targeting the AAVS1 locus downstream of the endogenous promoter/splice donor. Co-delivery of a promoter-less EGFP targeting vector in rAAV6 resulted in successful HR in 77.2 ± 3.0% of NK cells based on EGFP expression (**Figure 1H and 1I**). Together, these data demonstrate that high-efficiency Cas9-mediated gene KO and KI is achievable in activated primary human NK cells.

ADAM17 is responsible for the rapid cleavage of the activating Fc-gamma receptor CD16a from the surface of NK cells after activation^11,28^, resulting in inhibition of NK cytotoxicity. Chemical inhibitors of ADAM17 are currently in clinical trials in combination with antibody treatments as a method of enhancing the therapeutic effect of NK cells (NCT02141451). We reasoned that targeting ADAM17 directly in the NK cell could avoid systemic toxicities associated with off-target effects of chemical inhibitors. Using an artificial activation system (PMA)^11^ we demonstrate that ADAM17 KO NK cells maintain significantly higher surface expression of CD16a compared to control NK cells (which received Cas9 mRNA alone) and are on par with what is observed when NK cells are pre-treated with the ADAM17 inhibitor INCB007839 (**Figure 2A; Supplemental Figure 3A**). Surface membrane levels of CD62L, an additional target of ADAM17^11,29^, are also undiminished in activated ADAM17 KO NK cells (**Supplemental Figure 3B**). To test if ADAM17 KO NK cells have enhanced ADCC we performed standard ADCC assays using the CD20-positive Burkitt’s lymphoma cell line Raji. Raji cells were pre-treated with Rituximab, a monoclonal antibody targeting CD20. Rituximab-coated Raji cells induced cleavage of CD16a in 79.5 ± 1.3% of control NK cells, but there were no differences in CD16a cleavage in ADAM17 KO NK cells (**Figure 2B; Supplemental Figure 3C**). Moreover, while no significant difference was observed in cytotoxic degranulation (**Figure 2C; Supplemental Figure 3D**) ADAM17 KO NK cells displayed a significant increase in IFNγ production (**Figure 2D; Supplemental Figure 3E**). Importantly, ADCC was significantly enhanced when using ADAM17 KO versus control NK cells based on increased apoptosis as measured by cleaved Caspase 3 expression of Rituximab-coated Raji cells. (**Figure 2E; Supplemental Figure 3F)**. This finding contrasts with results using the clinically approved ADAM17 inhibitor, which did not lead to enhanced killing of Rituximab-coated Raji cells in previous reports^11^. Taken together, these data demonstrate that genetic KO of ADAM17 in NK cells blocks CD16a shedding comparable to ADAM17 inhibitors and leads to enhanced ADCC, not seen with current inhibitors of ADAM17.

**Figure 2.**
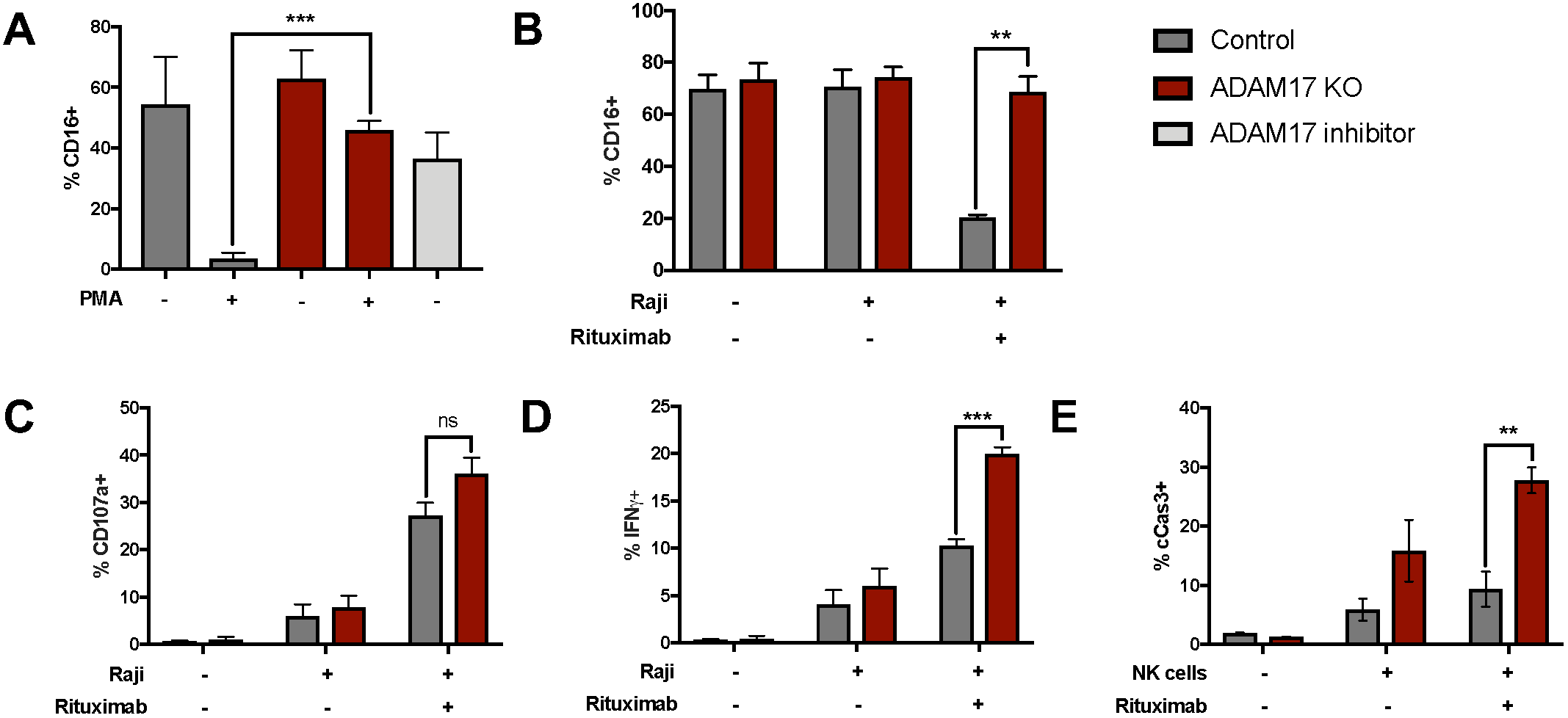
ADAM17 KO NK cells demonstrate enhanced ADCC. **(A)** Control (dark gray) and ADAM17 KO (maroon) NK cells were treated for one hour with 10 uM ADAM17 inhibitor (light gray) or DMSO. Cells were then stimulated with PMA (1 μg/mL) for one hour and CD16a expression was analyzed by flow cytometry (n=3 independent donors, ***P=0.0003, Student’s t test). **(B-E)** Raji cells were labeled for 10 minutes with CellTrace CFSE then incubated for 30 minutes with 10 ug/mL Rituximab. Raji cells were then co-cultured at a 2:1 (E:T) ratio with control (dark gray) or ADAM17 KO (maroon) NK cells for 6 hours (n=3 independent donors). **(B)** NK cell CD16A expression (**P=0.0012, Student’s t test), **(C)** NK cell degranulation, **(D)** NK cell IFNγ production (***P=0.0006, Student’s t test), **(E)** and Raji cell apoptosis (**P=0.007, Student’s t test).

NK cells are also capable of target cell killing independent of ADCC, through direct interaction between activating receptors on NK cells and cell surface proteins on target cells. However, tumor cells can block this cytotoxicity by inducing NK cell inhibitory signals through cell-surface protein interactions such as PD-L1:PD1^13,14^. We used our gene-editing method to generate PD1 KO NK cells and tested their ability to directly kill four cancer cell lines compared to control NK cells that were electroporated with Cas9 mRNA only. We selected four cancer cell lines with various expression levels of PD-L1 (**Supplemental Figure 4A**) – the chronic myeloid leukemia line K562 (PD-L1 low), the acute monocytic leukemia line THP1 (PD-L1 high), the prostate carcinoma line DU145 (PD-L1 high), and the ovarian carcinoma line MA148 (PD-L1 low). In a co-culture killing assay, PD1 KO NK cells had significantly higher levels of degranulation (**Figure 3A; Supplemental Figure 4B)** and production of IFNγ (**Figure 3B; Supplemental Figure 4C**). Similar to our ADCC results, PD1 KO NK cells also led to increased apoptosis of target cells compared to all controls (**Figure 3C; Supplemental Figure 4D**). To test functionality *in vivo*, we used PD1 KO NK cells to treat an orthotopic mouse xenograft model of ovarian cancer^30^. After xenograft establishment, mice were treated by a single intraperitoneal injection of rested control NK cells, donor-matched activated control NK cells which received Cas9 mRNA only, or donor-matched activated PD1 KO NK cells. Treatment with PD1 KO NK cells significantly reduced tumor burden (**Figure 3E**) and prolonged survival (**Figure 3F**) compared to both activated and rested NK cells, indicating improved therapeutic efficacy. To determine whether the differences we observed in NK cell effectiveness *in vivo* were due to increased PD-L1 expression by MA148 tumor cells after transplant, we harvested MA148 ascites tumors from mice treated with activated NK cells during early (week 1), middle (week 5), and late (week 10) timepoints after transplant. PD-L1 expression significantly increased over time after transplant and subsequent treatment with activated NK cells (**Figure 3G**). We also assessed PD-L1 expression on MA148 ascites tumors at the time of necropsy from mice in all treatment groups (**Figure 3H)** Overall, we observed significantly higher PD-L1 expression on MA148 cells from mice treated with activated NK cells. These data are consistent with the hypothesis that the inflammatory environment created by activated NK cells increases PD-L1 expression on tumor cells and underlies the need for creating PD1-deficient NK cells for cancer therapy.

**Figure 3.**
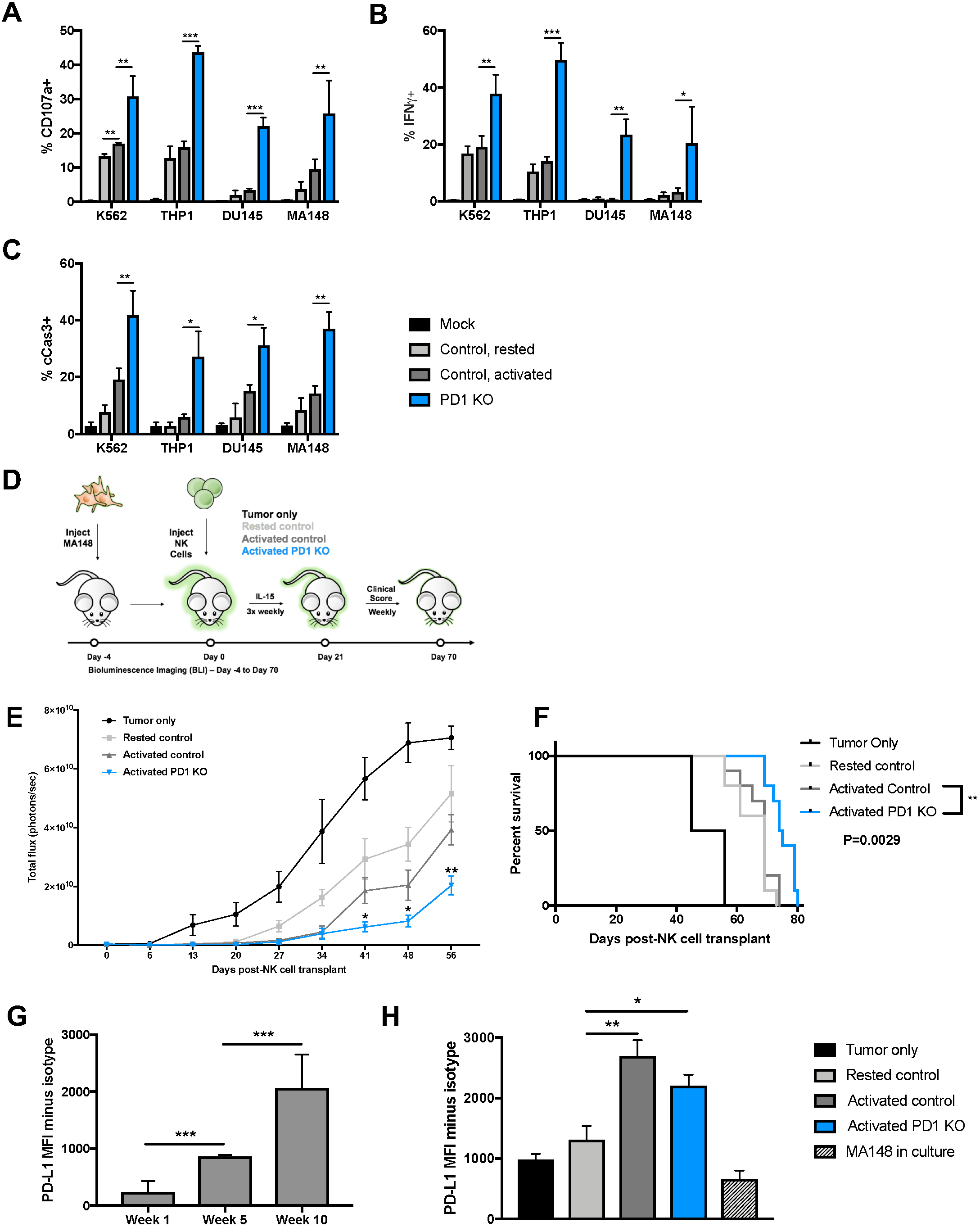
PD1 KO NK cells demonstrate enhanced anti-tumor function *in vitro* and *in vivo*. **(A-C)** Control NK cells were rested in 1 ng/mL IL15 (light gray) to support survival as previously described^35^. (light gray) Control (dark gray) and PD1 KO (blue) NK cells were activated for 1 week with Clone9.mbIL21 feeder cells. NK cells were then co-cultured at a 1:1 (E:T) ratio with target cell lines K562, THP1, DU145, and MA148 for 6 hours (n=3 independent donors, **P<0.01, ***P<0.001). **(A)** NK cell degranulation, **(B)** NK cell IFNγ production, and **(C)** target cell apoptosis. **(D-H)** 2.5×10^5^ luciferase-expressing MA148 cells were delivered IP to NSG mice. On day 4, PBS (black), or 1×10^6^ rested control (light gray), activated control (dark gray), or activated PD1 KO (blue) NK cells were delivered IP. NK cells were supported with a 5 ug dose of IL15 3 times weekly for 3 weeks. Tumor burden was assessed with bioluminescence imaging weekly. (n=8–10 mice per group, *P<0.05, **P<0.01, Student’s t test) **(E)** Tumor burden and **(F)** overall survival. **(G)** Mice treated with activated NK cells were sacrificed at week 1, week 5, and week 10 after treatment. MA148 cells were removed from IP ascites fluid and PD-L1 expression was measured by flow cytometry (n=4, ***P<0.001, Student’s t test). **(H)** MA148 cells were also harvested from IP ascites fluid from all treatment groups at sacrifice and PD-L1 expression was analyzed by flow cytometry (n=4, *P=0.0403, **P=0.0017, One-way ANOVA with Tukey post-hoc test).

CISH has been identified as a negative regulator of IL-15 signaling in murine NK cells, and its deletion has been shown to enhance their proliferation, activation, and tumor cell killing^25,31^. To determine whether CISH serves as a similar checkpoint in human NK cells, we performed assays for proliferation, activation, and tumor cell killing using CISH KO NK cells. In co-culture experiments with K562 or MA148 target cells, CISH KO NK cells did not have enhanced degranulation, cytokine production, or tumor cell killing (**Supplemental Figure 2A-C**). However, we did observe enhanced proliferation of CISH KO NK cells in response to low doses of IL-15 (**Supplemental Figure 2D and 2E**). These data emphasize the importance of validating results from murine NK cell experiments in human NK cells as significant differences have been previously reported between murine and human NK cells. These include expression of defining markers and signaling molecules^32^, localization to lymph nodes^33^, and cytotoxic capabilities^34^. Our data suggest that although CISH may be an important checkpoint in murine NK cells, it does not dampen antitumor immunity in human NK cells *in vitro*. Alternatively, the activated nature of our NK cells, due to our expansion culture conditions, could be superseding the normal regulatory effects of CISH in human NK cells.

NK cell immunotherapy holds great promise as an off-the-shelf cell therapy that does not require antigen specificity. Our work demonstrates the feasibility of generating clinical quantities of gene-edited NK cells with enhanced cytotoxic capabilities. We are able to effectively alter inhibitory protein expression in activated NK cells isolated from PBMCs. *In vitro* and *in vivo* assays demonstrate that these NK cells are capable of enhanced killing through ADCC and non-ADCC pathways. Future work will focus on methods of editing multiple genes simultaneously along with delivery of engineered receptors, similar to chimeric antigen receptors used in T cell therapy, for enhancing NK cell immunotherapy. If successful, these highly engineered, allogeneic NK cells could rival current autologous CAR-T cell therapies at a greatly reduced cost.

## References

1. Murphy, W. J., Parham, P. & Miller, J. S. NK Cells—From Bench to Clinic. Biol. Blood Marrow Transplant. 18, S2–S7 (2012).

2. Davis, Z. B., Felices, M., Verneris, M. R. & Miller, J. S. Natural Killer Cell Adoptive Transfer Therapy: Exploiting the First Line of Defense Against Cancer. Cancer J. 21, 486–491 (2015).

3. Davis, Z. B., Vallera, D. A., Miller, J. S. & Felices, M. Natural killer cells unleashed: Checkpoint receptor blockade and BiKE/TriKE utilization in NK-mediated anti-tumor immunotherapy. Semin. Immunol. 31, 64–75 (2017).

4. Podgorny, P. J. et al. Immune Cell Subset Counts Associated with Graft-versus-Host Disease. Biol. Blood Marrow Transplant. 20, 450–462 (2014).

5. Knorr, D. A., Bachanova, V., Verneris, M. R. & Miller, J. S. Clinical utility of natural killer cells in cancer therapy and transplantation. Semin. Immunol. 26, 161–172 (2014).

6. Schmiedel, D. et al. The RNA binding protein IMP3 facilitates tumor immune escape by downregulating the stress-induced ligands ULPB2 and MICB. eLife (2016). doi:10.7554/eLife.13426

7. Hofer, E. & Koehl, U. Natural Killer Cell-Based Cancer Immunotherapies: From Immune Evasion to Promising Targeted Cellular Therapies. Front. Immunol. 8, (2017).

8. Keating, G. M. Rituximab. Drugs 70, 1445–1476 (2010).

9. The impact of Fc-γ receptor polymorphisms in elderly patients with diffuse large B-cell lymphoma treated with CHOP with or without rituximab | Blood Journal. Available at: http://www.bloodjournal.org.ezp1.lib.umn.edu/content/118/17/4657?ijkey=f0f5581b19d99fd5c805459e07f2a3e2367a5327&keytype2=tf_ipsecsha. (Accessed: 23rd January 2018)

10. Mishra, H. K., Pore, N., Michelotti, E. F. & Walcheck, B. Anti-ADAM17 monoclonal antibody MEDI3622 increases IFNγ production by human NK cells in the presence of antibody-bound tumor cells. Cancer Immunol. Immunother. 67, 1407–1416 (2018).

11. Romee, R. et al. NK cell CD16 surface expression and function is regulated by a disintegrin and metalloprotease-17 (ADAM17). Blood 121, 3599–3608 (2013).

12. Effect of tumor cells and tumor microenvironment on NK-cell function - Vitale - 2014 - European Journal of Immunology - Wiley Online Library. Available at: http://onlinelibrary.wiley.com/doi/10.1002/eji.201344272/abstract;jsessionid=35BDC66F92538B0CA1758AFC15CDC67F.f04t03. (Accessed: 15th November 2017)

13. Guo, Y. et al. PD1 blockade enhances cytotoxicity of in vitro expanded natural killer cells towards myeloma cells. Oncotarget 7, 48360–48374 (2016).

14. Ray, A. et al. Targeting PD1-PDL1 immune checkpoint in plasmacytoid dendritic cell interactions with T cells, natural killer cells and multiple myeloma cells. Leukemia 29, 1441 (2015).

15. Vey, N. et al. A phase 1 trial of the anti-inhibitory KIR mAb IPH2101 for AML in complete remission. Blood 120, 4317–4323 (2012).

16. Chan, C. J. et al. The receptors CD96 and CD226 oppose each other in the regulation of natural killer cell functions. Nat. Immunol. 15, ni.2850 (2014).

17. Lesokhin, A. M., Callahan, M. K., Postow, M. A. & Wolchok, J. D. On being less tolerant: Enhanced cancer immunosurveillance enabled by targeting checkpoints and agonists of T cell activation. Sci. Transl. Med. 7, 280sr1–280sr1 (2015).

18. Denman, C. J. et al. Membrane-bound IL-21 promotes sustained ex vivo proliferation of human natural killer cells. PloS One 7, e30264 (2012).

19. Pierson, B. A., McGLAVE, P. B., Hu, W.-S. & Miller, J. S. Natural Killer Cell Proliferation Is Dependent on Human Serum and Markedly Increased Utilizing an Enriched Supplemented Basal Medium. J. Hematother. 4, 149–158 (1995).

20. Wang, X. et al. CRISPR-DAV: CRISPR NGS data analysis and visualization pipeline. Bioinforma. Oxf. Engl. 33, 3811–3812 (2017).

21. Osborn, M. J. et al. Evaluation of TCR Gene Editing Achieved by TALENs, CRISPR/Cas9, and megaTAL Nucleases. Mol. Ther. 24, 570–581 (2016).

22. Engineering of Primary Human B cells with CRISPR/Cas9 Targeted Nuclease | Scientific Reports. Available at: http://www.nature.com.ezp1.lib.umn.edu/articles/s41598-018-30358-0. (Accessed: 21st September 2018)

23. Shah, N. et al. Phase I study of cord blood-derived natural killer cells combined with autologous stem cell transplantation in multiple myeloma. Br. J. Haematol. 177, 457–466 (2017).

24. Hendel, A. et al. Chemically modified guide RNAs enhance CRISPR-Cas genome editing in human primary cells. Nat. Biotechnol. 33, nbt.3290 (2015).

25. Delconte, R. B. et al. CIS is a potent checkpoint in NK cell-mediated tumor immunity. Nat. Immunol. 17, 816 (2016).

26. Brinkman, E. K., Chen, T., Amendola, M. & van Steensel, B. Easy quantitative assessment of genome editing by sequence trace decomposition. Nucleic Acids Res. 42, e168–e168 (2014).

27. Lapteva, N., Szmania, S. M., van Rhee, F. & Rooney, C. M. Clinical Grade Purification and Expansion of Natural Killer Cells. Crit. Rev. Oncog. 19, 121–132 (2014).

28. Jing, Y. et al. Identification of an ADAM17 Cleavage Region in Human CD16 (FcγRIII) and the Engineering of a Non-Cleavable Version of the Receptor in NK Cells. PLOS ONE 10, e0121788 (2015).

29. ADAM17 deficiency by mature neutrophils has differential effects on L-selectin shedding | Blood Journal. Available at: http://www.bloodjournal.org.ezp1.lib.umn.edu/content/108/7/2275.long?sso-checked=true. (Accessed: 28th September 2018)

30. Felices, M. et al. IL-15 super-agonist (ALT-803) enhances natural killer (NK) cell function against ovarian cancer. Gynecol. Oncol. 145, 453–461 (2017).

31. Putz, E. M. et al. Targeting cytokine signaling checkpoint CIS activates NK cells to protect from tumor initiation and metastasis. OncoImmunology 6, e1267892 (2017).

32. Sungur, C. M. & Murphy, W. J. Utilization of mouse models to decipher natural killer cell biology and potential clinical applications. ASH Educ. Program Book 2013, 227–233 (2013).

33. Martín-Fontecha, A. et al. Induced recruitment of NK cells to lymph nodes provides IFN-γ for T_H_1 priming. Nat. Immunol. 5, 1260–1265 (2004).

34. Hébert, P. & Pruett, S. B. Selective Loss of Viability of Mouse NK Cells in Culture is Associated with Decreased NK Cell Lytic Function. In Vitr. Mol. Toxicol. 14, 71–82 (2001).

35. Romee, R. et al. Cytokine activation induces human memory-like NK cells. Blood 120, 4751–4760 (2012).

